# Intercalative DNA binding governs fluorescence enhancement of SYBR Gold

**DOI:** 10.1101/2020.05.23.112631

**Authors:** Pauline J. Kolbeck, Willem Vanderlinden, Thomas Nicolaus, Christian Gebhardt, Thorben Cordes, Jan Lipfert

## Abstract

SYBR Gold is a commonly used and particularly bright fluorescent DNA stain, however, its binding mode to DNA remains controversial. Here, we quantitate SYBR Gold binding to DNA using two complementary approaches. We use mechanical micromanipulation with magnetic tweezers (MT) to determine the effects of SYBR Gold binding on DNA length, twist, and mechanical properties. The MT assay reveals systematic lengthening and unwinding of DNA upon SYBR Gold binding, consistent with an intercalative binding mode where every SYBR Gold molecule unwinds DNA by 19.1° ± 0.7°. We complement the MT data with a spectroscopic characterization of SYBR Gold fluorescence upon addition to DNA. The data are well described by a global binding model for dye concentrations ≤1 μM, with binding parameters that quantitatively agree with the MT results. The fluorescence signal increases linearly with the number of intercalated SYBR Gold molecules. At dye concentrations >1 μM, fluorescence quenching and inner filter effects become relevant and it is required to correct the SYBR Gold fluorescence signals for quantitative assessment of DNA concentrations. In summary, we provide a mechanistic understanding of DNA-SYBR Gold interactions and present practical guidelines for optimal DNA detection and quantitative DNA sensing applications using SYBR Gold.

## INTRODUCTION

The interaction of DNA with ligands is fundamental for many cellular processes as well as biotechnological applications. In particular, fluorescent dyes are routinely used to label DNA for visualization and quantification in a wide variety of assays ranging from imaging of cells to analysis and quantification of gel bands or PCR products. SYBR Gold is a popular stain with very high sensitivity owing to the >1000-fold increase in fluorescence quantum yield on binding to DNA (1,2). Despite its widespread use, there is disagreement whether SYBR Gold binds in an intercalative (2,3) or in a minor-groove binding mode (4–6).

In general, the binding mode of DNA dyes impacts how binding depends on environmental conditions, DNA chain topology, or sequence context. Conversely, small-molecule binding to DNA can alter its structure and mechanical properties. Specifically, intercalation lengthens and unwinds the DNA helix (7,8), and the changes upon intercalation into DNA have been investigated at the single-molecule level using optical tweezers (3,9–21), AFM force spectroscopy (22,23), and magnetic tweezers (MT) (24–27). In contrast, minor groove binding has only much smaller effects, if any, on DNA length and winding angle (24).

Detection of DNA e.g. after separation by gel electrophoresis requires labeling and staining for visualization and quantitation. The extent of DNA binding depends on both on DNA and dye concentration, as with any equilibrium binding reaction. We need to quantitatively understand the binding properties of SYBR Gold to DNA in order to determine conditions for optimal signal-to-noise ratios when detecting DNA and conditions required to obtain a linear relationship between the amount of DNA and fluorescence intensity, which is desirable for quantitation.

To determine the binding mode of SYBR Gold to DNA and to investigate the fluorescence response under varying DNA and SYBR Gold concentrations, we combined single-molecule micromanipulation experiments with a range of fluorescence measurements. We present a binding model that describes both the single-molecule manipulation and bulk fluorescence data quantitatively. Binding parameters, i.e., the binding constant (dissociation constant) *K*_*d*_ and the binding site size *n*, were determined independently from both single-molecule manipulation and fluorescence experiments and were found to quantitatively agree. The close agreement between binding parameters determined from nano-manipulation and fluorescence strongly suggests that intercalation is the sole binding mode of SYBR Gold that enhances fluorescence upon DNA binding. In addition, we observe quenching of fluorescence at SYBR Gold concentrations > 1 μM and distinguish quenching mechanisms by fluorescence lifetimes measurements. Based on our experimental results, we present practical guidelines for optimal DNA detection and quantitative DNA sensing applications using SYBR Gold.

## MATERIALS & METHODS

### Chemicals

SYBR Gold in DMSO was purchased from Invitrogen. The concentration of the stock solution is not reported by the vendor, but was estimated from the absorbance at 495 nm assuming an extinction coefficient *ε*_495 nm_ = 75000 M^−1^cm^−1^ based on the spectral similarity of SYBR Gold and SYBR Green (2) (another SYBR family dye). Linear extrapolation from absorbance measurements on a serial dilution yields the concentration of the stock solution as [SYBR Gold] = (5.9 ± 0.1) mM (mean ± standard error).

### DNA constructs

For MT measurements, we use a 7.9-kbp DNA construct, prepared as described previously (24). The construct is generated by ligating handles (~600 bp) with either multiple biotin or multiple digoxigenin moieties fragments to an unmodified central DNA segment 7.9 kbp in length. Fluorescence intensity measurements of SYBR Gold are recorded in the presence of linear pBR322 plasmid DNA (NEB), which is produced by restriction of supercoiled circular pBR322 using restriction enzyme EcoRV (NEB) according to the protocol provided by the manufacturer. Completion of the linearization reaction was validated by agarose gel electrophoresis. Absorption, excitation and emission spectra of SYBR Gold are recorded in the presence of lambda phage DNA (NEB), which was dialyzed against phosphate buffered saline (PBS) prior to use.

### Magnetic tweezers setup

Experiments on DNA are performed on a home-built MT setup described previously (28). Two magnets (5 × 5 × 5 mm^3^; W-05-N50-G, Supermagnete) are placed in vertical configuration (29,30) on a motorized arm with a translational motor (M-126.PD2 motor with C-863.11-Mercury controller, Physik Instrumente) as well as a rotational motor (C-150.PD motor with C-863.11-Mercury controller, Physik Instrumente) to control the magnets’ rotation and z-position. The flow cell outlet is connected to a pump (ISM832C, Ismatec) for fluid handling. The setup is controlled using a Lab-VIEW software (National Instruments) described by Cnossen *et al.* (31).

Flow cells are built from two coverslips (24 × 60 mm, Carl Roth, Germany). The bottom coverslip was first functionalized using (3-Glycidoxypropyl)trimethoxysilane (abcr GmbH, Germany) and consecutively incubated with 50 μl of a 5000× diluted stock solution of polystyrene beads (Polysciences, USA) in ethanol (Carl Roth, Germany) to serve as reference beads for drift correction. The top coverslip has two openings with a radius of 1 mm for liquid exchange in the flow channel. The bottom and the top coverslip are glued together by a single layer of melted Parafilm (Carl Roth, Germany), precut to form a ~50 μl channel connecting the inlet and the outlet opening of the flow cell. After the flow cell assembly, one flow cell volume of 100 μg/ml anti-digoxigenin (Roche, Switzerland) in 1x PBS is introduced, and incubated overnight (at least 12 h). Subsequently, the flow cell is rinsed with 1 ml of 1x PBS and then passivated using a commercial passivation mix (BlockAid Blocking Solution, Thermoscientific) for 1 h to minimize non-specific interactions. Unbound material is removed from the flow cell by flushing with 1 ml of 1x PBS.

As magnetic beads we use 1 μm diameter MyOne beads (Life Technologies, USA). The DNA construct is attached to the streptavidin coated beads by incubating 0.5 μl of picomolar DNA stock solution and 2 μl beads in 250 μl 1x phosphate buffered saline (PBS, Sigma-Aldrich, USA) for 5 min. Then, the bead-coupled DNA constructs are introduced into the flow cell to bind to the flow cell surface via multiple digoxigenin:anti-digoxigenin bonds.

### Magnetic tweezers measurements

Prior to the experiments, DNA tethered beads are screened for the presence of multiple DNA tethers and torsional constraint by measuring their response to force and torque. To find out whether a magnetic bead is bound by more than one DNA tether to the surface, we introduce negative turns under high tension (*F* = 5 pN). In the case of a single double-stranded DNA tether, high tension prevents the formation of plectonemes at negative linking differences due to DNA melting and consequently, no change in height is observed. In contrast, if a bead is attached via two or more double-stranded DNA molecules, the molecules form braids when the bead is rotated causing a decrease in tether extension. Beads bound by multiple tethers are discarded from further analysis. To evaluate the presence of single strand breaks, positive linking differences are introduced at low force (F = 0.4 pN). Overwinding of torsionally constrained DNA leads to the formation of plectonemes, which decrease the z-extension, whereas in nicked DNA tethers, no linking difference can be induced, and the z-extension remains constant on magnet rotation.

For force-extension analysis, we exclusively examine torsionally unconstrained (nicked) DNA tethers. At first, we calibrate the magnet distance-to-force relation for each bead by recording the transverse fluctuations of the beads at different magnet separations for times approximately 100-fold larger than the characteristic time of the system at the corresponding force, and analyze the power spectral density of the fluctuations to quantify the force at each magnet position (32,33). The force-extension relation was then fitted using the worm-like chain (WLC) model (34) to extract the contour length and bending persistence length of the DNA. Next, we record the force-extension behavior in the presence of different concentrations of SYBR Gold. To this end, we first introduce 200 μl of the lowest dye dilution and measure the tether extension at 25 magnet positions between 0.1 mm and 7 mm. This experiment is repeated for 9 different SYBR Gold concentrations in increasing order. We use the previously calibrated force for each bead to construct force-extension curves which we subsequently fit using the WLC model to provide the contour length and persistence length as a function of SYBR Gold concentration.

Rotation curve measurements start by introducing 200 μl of 1x PBS in the flow cell using the peristaltic pump, at a flow rate of ~ 300 μl min^−1^. During flushing, the magnets are moved close to the flow cell to establish a force of 6.5 pN. The pulling force helps to prevent the magnetic beads from getting stuck on the surface, and, importantly, the field constrains the free rotation of the bead (35) during flushing, which is a requirement for determining the absolute shifts in DNA twist upon binding. During the actual measurement, the force is kept constant at 0.5 pN. While monitoring the DNA extension, we turn the magnet from negative to positive turns or from positive to negative turns (values vary for the different dye dilutions and are given in the result section). Then, we introduce 200 μl of the lowest SYBR Gold dilution and start another measurement again at a force of 0.5 pN. The experimental procedure is repeated for all SYBR Gold concentrations in increasing order. Further processing of the MT data was carried out using custom-written MATLAB routines.

### Absorption, excitation, and emission spectra

Emission spectra on excitation at 495 nm were recorded in the wavelength range 505-700 nm and excitation spectra on emission at 537 nm were recorded in the wavelength range 400-550 nm, employing a commercial spectrofluorometer (Fluoromax Plus; Horiba). The DNA concentration in the cuvette was stepwise decreased by replacing a fraction of the DNA solution with the same volume of a solution containing 1.2 μM SYBR Gold. Absorbance was recorded using an EvolutionTM 201/220 UV-Vis-spectrophotometer (ThermoFisher Scientific).

### Fluorescence intensity measurements

Linearized pBR322 plasmid DNA at varying concentrations was pipetted (25 μl) into a well of the well plate reader (Tecan Infinite M1000 PRO; Well plate: corning black polystyrene 384 well microplate with a flat bottom, Sigma-Aldrich, catalogue number: CLS3821). The fluorescence was read out from the bottom of the wells, with the excitation and emission bandwidth set to 5 nm, the gain to 100, the flash frequency to 400 Hz, and the integration time to 20 μs. We choose the excitation and emission wavelengths to be 495 nm and 537 nm (excitation and emission maxima for SYBR Gold, as provided by Invitrogen). For control measurements, lambda phage DNA (NEB) at a constant concentration of 3.5 ng/μl and varying SBR Gold concentration were used and performed otherwise identically.

For the qPCR cycler (CFX96 Touch Real-Time PCR Detection System, Bio Rad) fluorescence read out experiments, again linearized pBR322 DNA was used and dilution series were filled into low-profile PCR tubes (Bio Rad, product ID: TLS-0851), which were closed with flat, optical, ultra-clear caps (Bio Rad, product ID: TCS-0803). We used the channel with absorption and emission wavelengths of 494 nm and 518 nm, respectively, which are the closest match to those of SYBR Gold (495 nm and 537 nm, respectively) and read out the fluorescence intensities at 24 °C from the top.

For the gel electrophoresis we added Gel Loading Dye Purple (6x) (NEB) to pBR322 DNA. We used 1%-agarose (Carl Roth) gels and TAE buffer (40 mM Tris, 20 mM acetic acid, and 1 mM EDTA, pH 8.6). The gels were run for 120 min at 75 V. Afterwards the gel was removed from the gel box and placed for 20 minutes in 100 ml of 0.6, 1.5, or 3 μM SYBR Gold in TAE buffer, respectively, for staining. Subsequently, the gel was de-stained in TAE buffer for 15 min at room temperature. The gels were then visualized using a Gel Doc XR+ System (Biorad).

### Fluorescence lifetime measurements

Fluorescence lifetime measurements used a homebuilt apparatus for time-correlated single-photon counting (TCSPC). Pulsed excitation was available at 485 nm (PicoQuant LDH–D–C–485, controller: PicoQuant PDL 828 “Sepia II”, 20 or 13.33 MHz laser repetition rate) to excite the sample cuvette. A lambda-half waveplate (Laser Components ACWP-450-650-10-2-R30 AR/AR, Ø1”) and a linear polarizer (Edmund Optics POLARIZER 25.4mm DIA UNMTD) were used to rotate and adjust the excitation polarization to vertically polarized light. Light emitted by the sample after excitation was filtered for polarization with a second linear polarizer (Edmund Optics ULTRA BB WIREGRID POL 25RND MTD) under magic angle condition (54.7°) with respect to the vertical axis. The emission polarizer was mounted inside a 3D custom-printed chassis housing with automated rotation mount (Thorlabs CRM1/M, Reichelt GRABIT SERVO), which was controlled by a processing unit (Arduino Mega 2560) for automated rotation of the emission polarizer. A lens (Thorlabs AC254-110-A) was used for higher collection efficiencies in combination with filters (AHF 488 long pass filter BLP01-488-R-25 and notch filter ZET488NF) for removal of scattered laser light before the avalanche photodiode used for detection (Excelitas SPCM-AQRH-34). Data were recorded by a TCSPC unit (PicoQuant HydraHarp 400, 16 ps TCSPC time resolution) with a commercial control and evaluation software provided by the supplier (PicoQuant SymPho Time 64).

We diluted lambda phage DNA (NEB), dialyzed overnight against PBS, and SYBR Gold to the specified dye concentrations (0.03 to 60 μM) and analyzed the sample in a 10 × 2 mm^2^ cuvette (Perkin Elmer UV/VIS Spectroscopy Cell B0631122) for 5 min for each condition. The excitation laser power was 1.5 or 15 μW in order to operate in an optimal range with the photon detection rates between 20 and 80 kHz. Additionally, we measured the instrument response function (IRF) by replacing the cuvette with a silver mirror to direct the laser beam directly onto the APD chip.

For the fluorescence lifetime evaluation, we used the n-exponential reconvolution fit that is implemented in the measurement software SymPho Time 64. The optimization uses a maximum likelihood estimator, where the fluorescent signal *I*(*t*) is described as the convolution of the IRF (*t* − *t*_*off*_ with a triple exponential decay (36):

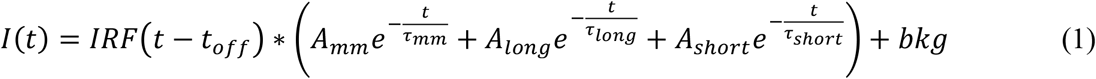

In this formula, the first exponential component 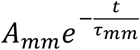 accounts for a small mismatch between measured IRF (with a mirror) and the “real” IRF in the cuvette measurements. This mismatch was compensated by including a fast decay component in the fitting procedure, with a fixed decay time *τ*_*mm*_ = 50 ps. The fluorophore lifetime was described with two exponential decays with lifetimes *τ*_*long*_ and *τ*_*short*_, accounting for a fast component, which becomes relevant at high dye concentrations, in addition to the dominant slow lifetime. The reported lifetimes in the main text are amplitude-weighted averages 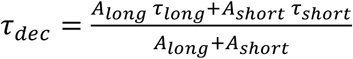 of the two lifetime components *τ*_*long*_ and *τ*_*short*_. The errors for *τ*_*dec*_ are derived from the fit uncertainty of all parameter based on standard error propagation rules for non-independent combined quantities.

### Binding models: McGhee-von Hippel model and DNA concentration effects

In order to describe the binding of SYBR Gold to double-stranded DNA in our single-molecule tweezers assays, where the DNA concentration is very small and, therefore, the free and total ligand concentrations approximately equal, we used the McGhee-von Hippel model of ligand-substrate binding (37) for the fractional number of molecules bound per base pair, γ:

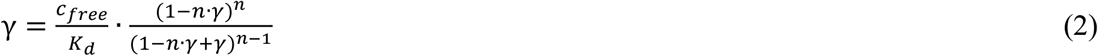

where *c*_*free*_ (≈ *c*_*total*_ under these conditions) is the free ligand concentration, *K*_*d*_ is the binding (dissociation) constant (in M) and *n* is the binding site size (in base pairs). γ was determined from the DNA contour length change with increasing ligand concentration determined in stretching experiments on rotationally unconstrained DNA as

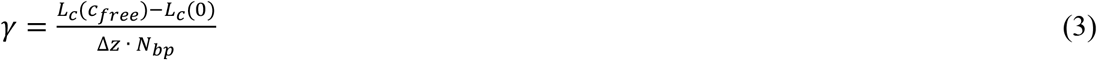

where *L*_*c*_(*c*_*free*_) is the DNA contour length at a certain ligand concentration, Δ*z* is the increase in DNA contour length per ligand bound, and *N*_*bp*_ is the number of base pairs of the DNA (*N*_*bp*_ = 7922 for our DNA constructs). We used a fixed value for the DNA contour length increase per SYBR Gold binding event Δ*z* = 0.34 nm, as was suggested previously for intercalators (15,24,38,39).

From bulk experiments using a plate reader, qPCR cycler, or a gel imager we determined the fluorescence intensity *I* as a function of DNA concentration *c*_*DNA*_ and dye concentrations *c*_*total*_. For bulk measurements, the assumption that *c*_*free*_ ≈ *c*_*total*_ often does not hold, as the free ligand concentration is on the order of the bound ligand concentration and both the finite dye and DNA concentrations need to be taken into account. Therefore, we rewrote Eqn. (2) in terms of *c*_*total*_, which is experimentally known, by expressing the total ligand concentration as the sum of free ligand concentration and bound ligand concentration:

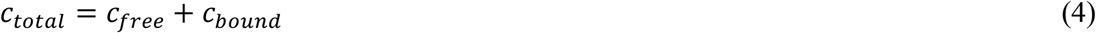

We define

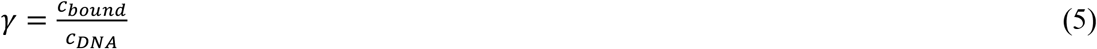

with *c*_*DNA*_ being the total DNA concentration which is also experimentally known. Subsequently, we combined Eqn. (4) and (5) and inserted them into Eqn. (2) to obtain:

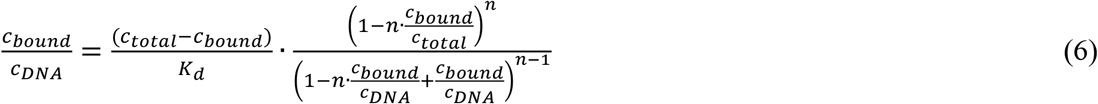

We call Eqn. (6) the McGhee-von Hippel model for finite DNA concentrations.

To fit our fluorescent intensity data with the McGhee-von Hippel model for finite DNA concentrations, we assume that the fluorescent intensity is proportional to the concentration of bound SYBR Gold molecules

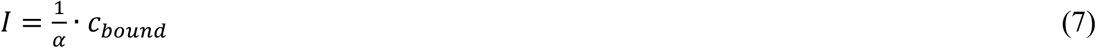

where *α* is a proportionality constant that we treated as a fitting parameter. It is reasonable to neglect fluorescence contributions from the free dye, as for SYBR Gold the fluorescent intensity increases by more than a factor of 1000 upon binding to double-stranded DNA (1,2). We substituted the concentration of bound SYBR Gold molecules *c*_*bound*_ in Eqn. (6) by *α* · *I* to obtain the final equation that can be fit to the fluorescence intensity data:

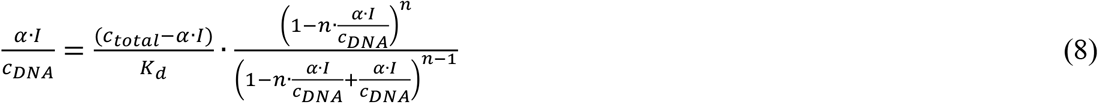

In Eqn. (8), the total ligand concentration *c*_*total*_ as well as the total DNA concentration *c*_*DNA*_ are experimentally known, the fluorescence intensity *I* is measured, and the dissociation constant *K*_*d*_, the binding site size *n*, and the proportionality factor *α* are fitting parameters. For the gel imager data, we need to take into account that the concentration in the staining solution is not equal to the final concentration in the gel, due to dilution by the gel volume, incomplete penetration of the dye into the gel, and the final de-staining step. We, therefore, rescale the SYBR Gold concentrations for all gel data with a single global dilution factor, which we determine to be ≈ 0.1.

## RESULTS

To investigate the binding mode of SYBR Gold to DNA and to quantitatively monitor the resulting changes in DNA properties and fluorescence, we use different, complementary techniques. In a first series of experiments, we use MT micromanipulation to monitor binding and examine the effects of the dye on the structural and mechanical properties of DNA under controlled stretching forces and degrees of supercoiling. In a second set of experiments, we monitor SYBR Gold fluorescence in the presence of various concentrations of dye and DNA via fluorescence spectroscopy. The MT experiments are performed at varying SYBR Gold concentrations using a custom-build multiplexed MT set up (**Fig 1A** and Materials and Methods).

**Figure 1.**
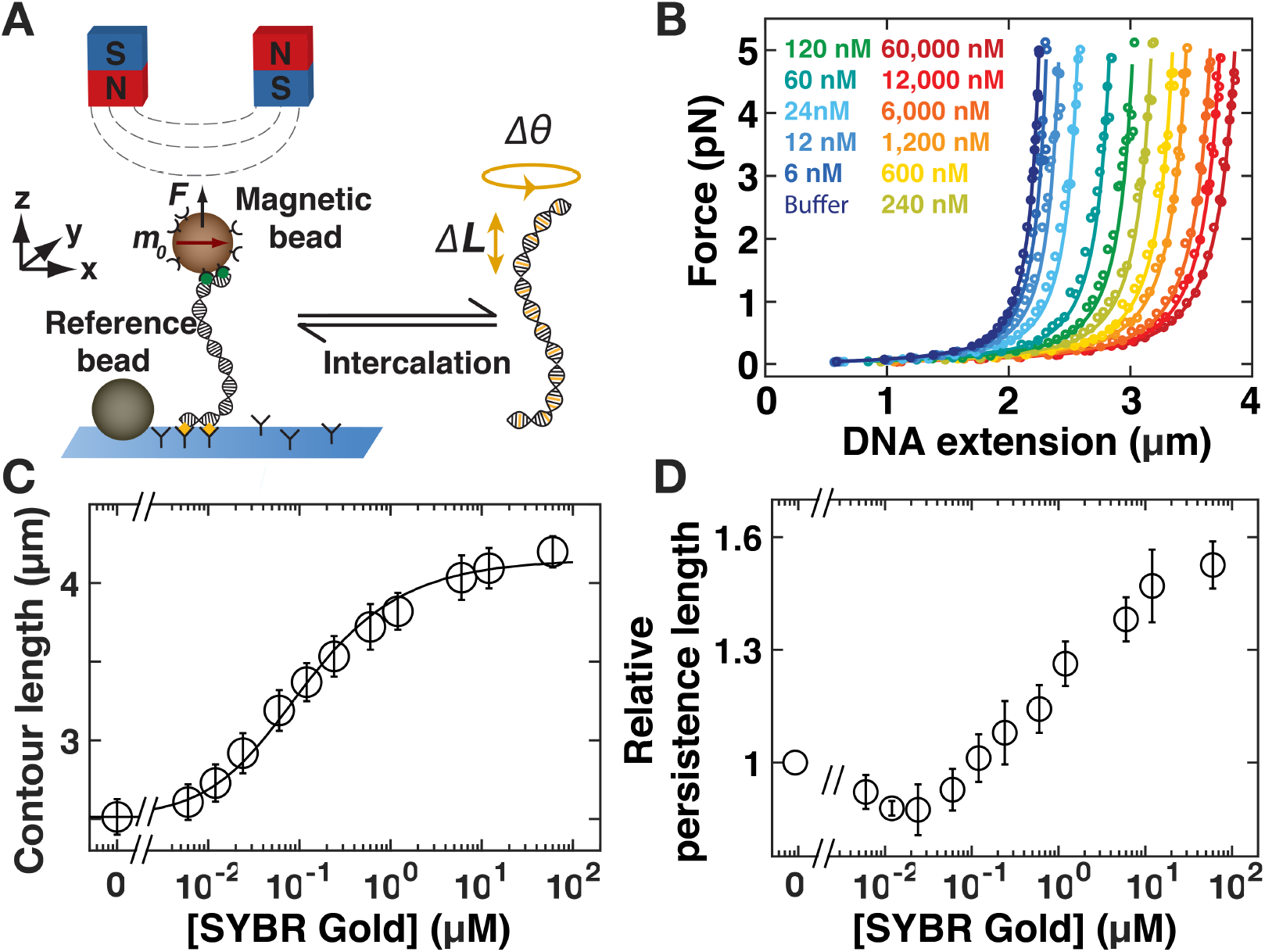
Effects of SYBR Gold on the force-extension behavior of DNA. **A)** Schematic of magnetic tweezers. DNA molecules are tethered between a surface and magnetic beads in a flow cell. Permanent magnets above the flow cell enable the application of stretching forces and torques, respectively. Upon intercalation, the DNA molecules lengthen and change their equilibrium twist. **B)** Force-extension curves for 7.9-kb DNA in the presence of increasing concentrations of SYBR Gold (increasing concentrations from blue to red indicated in the figure legend). Symbols are raw data, lines are fits of the WLC model. A systematic increase the DNA extension with increasing SYBR Gold concentration is apparent. **C)** DNA contour length determined from fits of the WLC model as a function of SYBR Gold concentration. The black line is a fit to the McGhee–von Hippel model (reduced *χ*^2^ = 0.18; see main text for details), with a dissociation constant *K*_*d*_ = 0.13 ± 0.01 μM and a binding site size n = 1.67 ± 0.04. **D)** DNA bending persistence length from WLC fits measured as a function of the dye concentration, normalized to the bending persistence length measured for bare DNA, indicating that the persistence length increases with increasing amount of SYBR Gold. Data points and error bars in panels C and D are the mean and standard deviation from at least ten independent measurements. In panel B one typical experiment is shown for clarity.

### SYBR Gold lengthens the DNA contour

To assess whether SYBR Gold binding lengthens the DNA contour, we first performed DNA stretching experiments in the presence of increasing concentrations of SYBR Gold on torsionally relaxed (nicked) DNA. We focused on the force regime below 5 pN for our force–extension measurements: in this regime the DNA extension is well described by the worm-like chain (WLC) model of entropic stretching elasticity (40) while enthalpic contributions to stretching and the force-dependence of intercalative binding are negligible (15,39). In the absence of SYBR Gold the response of dsDNA to force shows the characteristic response of entropic stretching elasticity (**Fig 1B,** dark blue circles). A fit to the WLC model (34) (**Fig 1B**, dark blue line) yields a contour length *L*_*C*_ = (2.4 ± 0.1) μm (mean and standard deviation from ten independent measurements), in close agreement with the expected crystallographic length of 2.6 μm expected for a 7.9-kb dsDNA. We then examined the force-extension response of torsionally unconstrained DNA at increasing concentrations of SYBR Gold (**Fig 1B**, curves from light blue to red). Fitting the WLC model to the force-extension data demonstrates that the contour length of the molecule systematically increases with increasing SYBR Gold concentration: compared to the contour length of bare DNA, the contour length of DNA in the presence of 6 μM SYBR Gold is increased by a factor of 1.67. A significant contour length increase upon binding by ~1.7 fold is consistent with an intercalative binding mode (9,15,24) and very unlikely to result from minor or major groove binding. The increase in the contour length follows a binding curve behavior and saturates at concentrations ≥ 1 μM (**Fig 1C)**. To quantitatively determine SYBR Gold binding parameters from the mechanical response of DNA, we employ the McGhee-von Hippel model (Equation 1 in Materials and Methods) (37). The fractional number of bound SYBR Gold molecules is computed from the fitted contour length *L*_*C*_(*c*) at a given SYBR Gold concentration *c* by Equation 2. We use a value for the length increase per SYBR Gold bound of Δ*z* = 0.34 nm, typical for intercalation (15,24,38). Fitting the McGhee-von Hippel model to the contour length data determined from our force-extension measurements, we find the dissociation constant *K*_*d*_ = (1.32 ± 0.13) · 10^−7^ M and a binding site size *n* = 1.67 ± 0.04. Errors are obtained from a “bootstrapping” procedure (41) by generating 1000 synthetic data sets from the experimental data and computing the standard deviation over repeated McGhee-von Hippel fits. The value of the binding site size *n* can be interpreted as SYBR Gold intercalation at saturation occurring slightly more than at every other basepair (42). In addition to an increase in DNA contour length upon SYBR Gold binding, the bending rigidity is also changing (**Fig 1D)**. We found *L*_*P*_= (35 ± 4) nm in the absence of SYBR Gold. This value is in agreement though slightly lower than reported data for DNA persistence length in PBS (24). With increasing dye concentration, the persistence length initially stays constant or decreases very lightly up to a SYBR Gold concentration of ~0.1 μM and then significantly increases at higher dye concentrations (**Fig 1D**). In comparison to the persistence length of bare DNA, the persistence length of DNA in the presence of 6 μM SYBR Gold increases by a factor of 1.55. Previous studies of intercalators have reported both decreases in bending stiffness, e.g. for ethidium bromide, PicoGreen, or a Ru(II) metallo-intercalator (24,43,44), but also constant or increasing stiffness, e.g. for the bis-intercalators YOYO and TOTO (26,45,46).

### SYBR Gold untwists DNA

Force-extension measurements of torsionally unconstrained DNA in the presence of different concentrations strongly suggest that SYBR Gold binds DNA in an intercalative binding mode. To further investigate the SYBR Gold binding to DNA, we probe the effects of varying concentrations of SYBR Gold on DNA twist using rotationally constrained DNA molecules. We control the DNA linking number *Lk* by rotation of the magnets and monitor DNA extension as a function of applied magnet turns at a low constant force *F* = 0.5 pN (**Fig 2A**). In the absence of SYBR Gold, we observe the characteristic response of bare DNA (**Fig 2A,** dark blue data): Initially the change in turns (*ΔLk*) leads to elastic twist deformations of DNA that minimally affect the tether extension. Subsequently, the DNA buckles to form plectonemic supercoils. In the plectonemic regime, a further increase of *ΔLk* results in a linear reduction of its end-to-end extension. For measurements at low forces (here *F* = 0.5 pN), the response of the DNA is symmetric about *Lk*_0_(i.e. zero applied turns, corresponding to a torsionally relaxed molecule): if we introduce positive or negative turns the DNA forms positive or negative plectonemic supercoils, respectively. Throughout, number of applied turns where the DNA tether was torsionally relaxed in the absence of added SYBR Gold, corresponding to *Lk* = *Lk*_0_ and determined as the midpoint of the symmetric rotation curve (**Supplementary Figure S1A**), is used as a reference. Addition of increasing concentration of SYBR Gold leads to dramatic changes in the shape and position of the extension vs. applied turns curves (**Figure 2A**, increasing dye concentration from blue to red). There are four effects that become increasingly pronounced with increasing dye concentration: i) an overall increase of the DNA tether length, ii) a shift of the centers of the curves towards negative turns, iii) a broadening of the curves, *i.e.* an extension of the pre-buckling regime, and iv) a negative slope of extension vs. turns in the pre-buckling regime. In the following, we will discuss each of these observations:

i. The increase in the extension with increasing SYBR Gold concentration is readily understood from the force-extension measurements discussed in the previous section. The center of the rotation-extension curves corresponds to torsionally relaxed DNA for which a systematic length increase with increasing SYBR Gold concentration by up to ~1.7-fold at saturating SYBR Gold was observed in the force-extension measurements (**Figure 1C**).
ii. The shift in center position of the rotation-extension curves with increasing SYBR Gold concentration compared to bare DNA is indicative of DNA unwinding upon SYBR Gold binding, which is again consistent with intercalation. We can understand the shift following the addition of SYBR Gold by considering that our DNA molecules are torsionally constrained (*Lk*; is a topological constant). If the binding of SYBR Gold causes a change in the DNA *Tw*, compensatory changes in *Wr* must occur, which, in turn, result in a reduction of the DNA end-to-end extension due to the formation of plectonemes. We quantified the shift in the extension curves by linearly fitting the extension vs. applied turns response in the plectonemic regime for both positive and negative plectonemes and determine the center of the curve as the intersection of the two slopes. Plotting the shift in the rotation curve centers as a function of the SYBR Gold concentration (**Fig 2B**), we again obtain a binding curve behavior. We fit the center shifts by *ΔTw*([*SYBR Gold*]) = *γ* · *Nbp* · *Δθ*, where we use *γ* as computed from the force extension measurements (**Figure 1**). The change in DNA twist per SYBR Gold binding event *Δθ* is treated as a fitting parameter. The resulting fit (**Fig 2B,** solid line) gives *Δθ* = 19.1° ± 0.7°. The error was computed as the standard deviation over fits to 1000 synthetic bootstrap data sets. We note that the last data point falls somewhat below the fitted curve, qualitatively similar to observations for ethidium bromide (24,47), which might indicate a tendency for cooperative binding at very high concentrations (47). However, we felt that the fit is still sufficiently accurate and does not warrant introduction of an additional parameter. The fitted value for *Δθ* is comparable to values found for other intercalators, for example 27.3° ± 1° for ethidium bromide (24,47) or 21° ± 14° for Pico Green (43). The slopes in the plectonemic regime *q* at *F* = 0.5 pN are the same, within experimental error, for positive and negative supercoils and also do not change, within error, with increasing concentrations of SYBR Gold (**Supplementary Figure S1B**). In contrast, for ethidium bromide the slopes decrease with increasing intercalator concentration (24). The slope in the plectonemic regime *q* is essentially determined by the size of the plectonemes (48–50). The size of the plectonemes and thus *q* increases with increasing bending persistence length (a simple model (49,51) suggests *q*~*L*_*P*_^1/2^) and increases with increasing charge density of the DNA (50) (approximately as *q* ~ rescaled charge density). For ethidium bromide, both the bending persistence length (24) and the charge density decrease through intercalation of the positively charged dye; the effects combine to give a significant decrease in *q* with increasing ethidium concentration. For SYBR Gold, the bending persistence length increases (**Figure 1D**) while the charge density decreases with increasing dye concentration; the effects appear to approximately cancel, such that the overall change in *q* is small.

**Figure 2.**
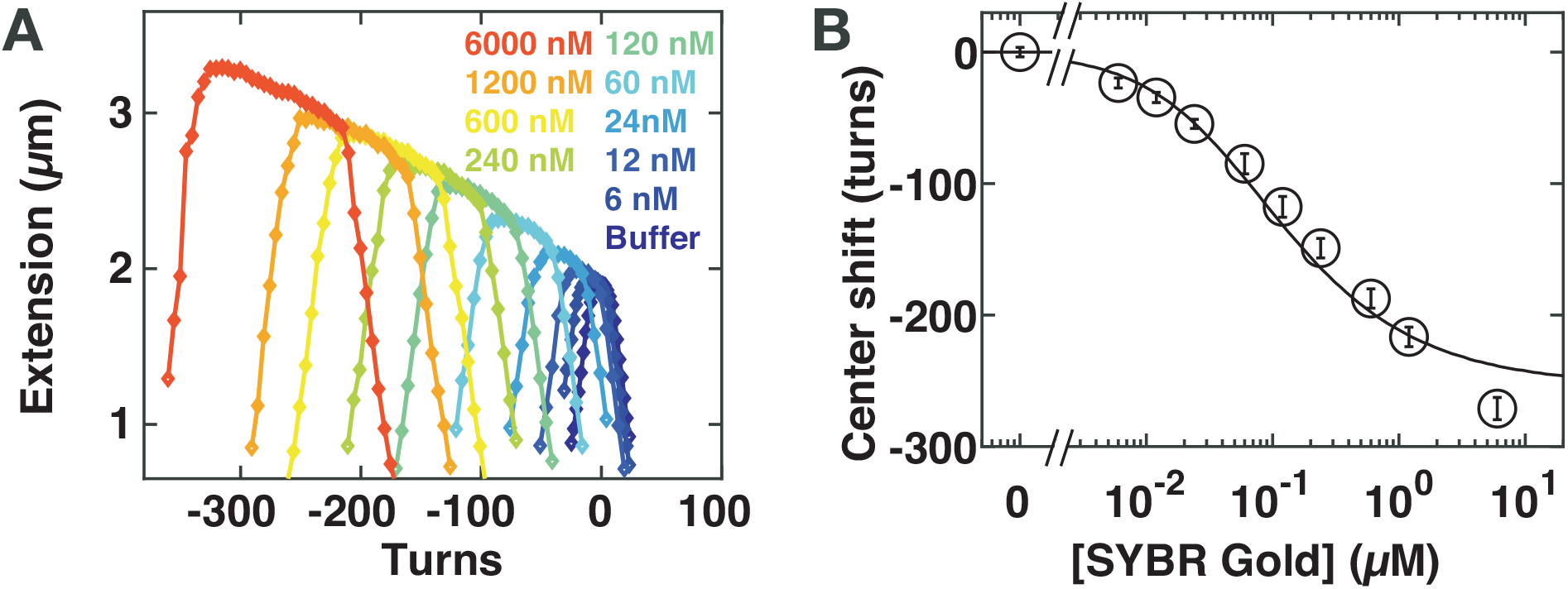
Effect of SYBR Gold on DNA twist and torque. **A)** Rotation-extension curves for 7.9 kb DNA at *F* = 0.5 pN in the presence of increasing concentrations of SYBR Gold. The SYBR Gold concentrations are (from blue to red) 0, 12, 24, 60, 120, 240, 600, 1200, 6000 nM. With increasing concentrations of SYBR Gold the rotation curves shift to negative turns; the DNA length at the center of the curves increases; and the rotation-extension curves broaden. **B)** Quantification of the shift in the center position of the rotation-extension curves as a function of the SYBR Gold concentration. The center positions were determined from fitting slopes in the positive and negative plectonemic regime and by computing the intersection of the two slopes. The black line is a fit of the McGhee-von Hippel model (reduced *χ*^2^ = 2.9; see main text for details), with the dissociation constant *K*_*d*_ and binding site size *n* set to the values determined from the force extension data (**Figure 1**) and the unwinding angle per SYBR Gold intercalation determined from the fit to be *Δθ* =19.1° ± 0.7°. Data points and error bars in panel B are the mean and standard deviation from at least 14 independent measurements. In panel A one typical experiment is shown for clarity.

(iii) and (iv) Broadening of the rotation curves and a slope in the pre-buckling regime (**Figure 2A** and **Supplementary Figure S1C**) are similar to what has been observed for other intercalators, notably ethidium bromide (24,47), PicoGreen (43), and the bis-intercalator YOYO-1 (27). The two effects can be understood from the properties of the DNA tethers and how they are changed upon SYBR Gold intercalation. Fundamentally, a molecule buckles if the energy required to form a plectoneme becomes less than the twist energy stored in the chain induced by adding turns to the molecule. For naked DNA in the pre-buckling regime, the torque builds up as *Γ* = (*k*_*B*_*T* · *C*/*L*_*C*_) · *2π* · *ΔLk*, where *k*_*B*_ is the Boltzmann constant, *T* the absolute temperature, and *C* the torsional persistence length. Buckling occurs once the built-up torque reaches the critical torque for buckling *Γ*_*buck*_, which increases with increasing bending persistence length, approximately (49,51) as *Γ*_*buck*_ = (*k*_*B*_*T* · *L*_*P*_ · *F*)^1/2^. Part of the broadening of the rotation curves can, therefore, likely be explained by changes in the mechanical properties of DNA in the presence of SYBR Gold. The observed increase in bending persistence length (**Figure 1D**) will tend to increase the number of turns required for buckling (yet only by 1.6^1/2^ ≈ 1.25-fold); similarly, a decrease in the torsional persistence length, as has been observed for ethidium bromide upon intercalation (52), would increase the number of turns required for buckling and broaden the curves (24).

However, in addition to changing mechanical properties in the presence of intercalation, torque-dependent intercalation will also contribute to the broadening of the rotation curves. The extension is maximal not at the center of the rotation-extension curve, as is the case for bare DNA, but at negative turns just before buckling, due to the overall slope in the pre-buckling regime. The dependence of tether length on the number of applied turns at a given force and SYBR Gold concentration suggests that the application of a negative torque increases SYBR Gold binding, whereas the application of a positive torque hinders SYBR Gold binding, which is to be expected from Le Chatelier’s principle as SYBR Gold intercalation unwinds the DNA helix. We note that the slopes of the plateaus in the pre-buckling regime corresponded roughly to the slope of the curve connecting the center position of the rotation-extension curves, similar to what has been observed for ethidium bromide (47). The center positions of the rotation-extension curves are given by the coupling between DNA elongation and untwisting upon intercalation. The observation that the slope in the pre-buckling plateaus matches the slope of the line connecting the rotation curve centers suggests that upon twisting SYBR Gold-bound DNA, the applied turns are predominantly absorbed by torque-induced intercalation, again suggesting an important role of torque-dependent binding, due to the unwinding of the helix upon intercalation. In conclusion, the results of our DNA micromanipulation experiments reveal that SYBR Gold binding to DNA systematically lengthens, by up to ~1.7-fold, and unwinds DNA, by 19.1° ± 0.7° per binding event, fully consistent with an intercalative binding mode.

### SYBR Gold absorption, excitation, and emission spectra

To relate the SYBR Gold binding behavior revealed by the MT measurements to SYBR Gold fluorescence, we first determined absorption, excitation, and emission spectra (**Figure 3**) of the dye in the presence of DNA. The absorption spectra show a systematic increase of absorbance with SYBR Gold concentration, at fixed DNA concentration, with a peak around 490 nm (**Figure 3A**) and a secondary peak around 300 nm, consistent with previous measurements (1,2). The position of the absorbance peak shifts to shorter wavelength with increasing SYBR Gold concentrations (**Figure 3C**). The position of the peak is well described by a model (**Figure 3C**, solid line) that assumes fixed absorbance peak wavelengths λ_max,free_ and λ_max,bound_ for free and bound SYBR Gold and a linear superposition of the form

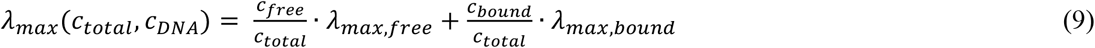

**Figure 3.**
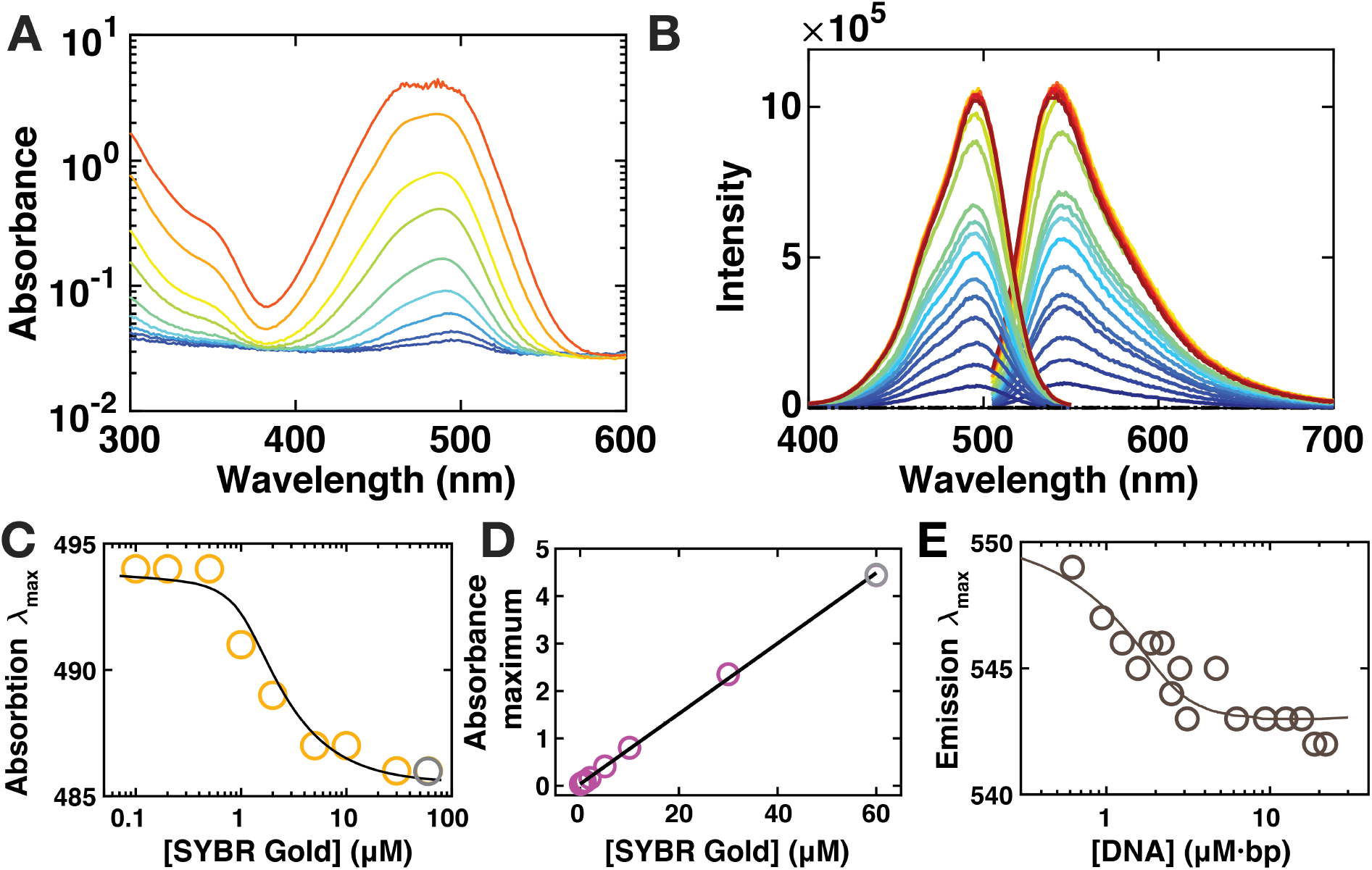
SYBR Gold absorbance, excitation, and emission spectra. **A)** Absorbance spectra for varying SYBR Gold concentrations (from blue to red: 0.1, 0.2, 0.5, 1, 2, 5, 10, 30, 60 μM) in the presence of 2 μM·bp DNA (Lambda-DNA (NEB)). The peak of the spectrum at the highest concentration is noisy, as the dynamic range of the instrument is approached. Points from this spectrum are included below in panel C and D, but greyed out. **B)** SYBR Gold excitation and emission spectra at constant SYBR Gold concentration (1.2 μM) and varying DNA concentration (from blue to red: 0.9, 1.3, 1.6, 1.9, 2.2, 2.8, 3.1, 4.7, 6.3, 9.4, 12.5, 15.6, 18.8, 21.9, 25.0, 28.1, 31.3 μM·bp, dotted line: SYBR Gold only, dashed line: PBS only. **C)** Position of the absorbance maxima of the SYBR Gold absorbance spectra as a function of the SYBR Gold concentration (circles). The solid line is a fit to the finite concentration McGhee-von Hippel model with the wavelengths for maximum absorbance of free λ_max,free_ and intercalated λ_max,bound_ SYBR Gold as the only fitting parameter (see main text). From the fit we find λ_max,free_ = 486 nm and λ_max,intercalated_ = 494 nm. **D)** Value of the absorbance maxima of the SYBR Gold absorbance spectra as a function of the SYBR Gold concentration. The circles are the experimental data, the solid line is a linear fit (R = 0.9997); extrapolation of the linear fit yields the value for the undiluted SYBR Gold which is found to be 435±6 and implies a concentration of the undiluted SYBR Gold of 5.9±0.1 mM. **E)** Position of the emission maxima at constant SYBR Gold concentration (1.2 μM) and varying DNA concentrations. The data are well described by the model in Equation 9, with binding parameters *K*_*d*_ and *n* fixed to the values determined from the MT measurements (analogous to panel C). From the fit we find for λ_max,free_ = 550 nm and λ_max,bound_ = 543 nm for emission.

The free and bound SYBR gold concentrations are computed from the finite concentration McGhee-von Hippel model (Equation 5), using the *K*_*d*_ and *n* values determined from the MT force-extension measurements (**Figure 1**). The absorbance peak wavelengths are determined from a fit as λ_max,free_ = 486 nm and λ_max,bound_ = 494 nm, very close to the previously reported values of 487 and 495 or 496 nm for free and bound SYBR Gold, respectively (1,2). The absorbance increases linearly with increasing SYBR Gold concentration (**Figure 3D**), which suggests that absorption does not depend on the dye being intercalated or in free solution (since the DNA concentration is fixed for the data set in Figure 3A). The slope of the linear dependence agrees well with the value for ε used in the concentration determination using a different instrument (**Figure 3D** and Methods), providing a consistency check.

The excitation and emission spectra exhibit single peaks in the range probed (**Figure 3B**). While the position of the excitation peak at 496 nm does not shift significantly upon binding (**Supplementary Figure S2**), we observe a systematic shift for the single emission peak at ~545 nm (**Figure 3E**). The emission data are well described by the model that was fit to the absorbance data (Equation 9), using again the finite concentration McGhee-von Hippel model with *K*_*d*_ and *n* fixed to the values determined from the MT measurements and fitting λ_max,free_ and λ_max,bound_ for emission. We find λ_max,free_ = 550 nm and λ_max,intercalated_ = 543 nm for emission, similar to, but slightly higher than the values reported previously (1,2). The difference is likely within experimental error and we note that the magnitude of the shift is very similar to what was previously reported. The fact that the shifts in the absorbance and emission spectra follow the same binding curve as determine from mechanical manipulation in the MT strongly suggests that intercalation is the relevant binding mode for fluorescence.

### Fluorescence intensity scales linearly with intercalated SYBR Gold at low concentrations

To further correlate our findings from DNA micromanipulation with SYBR Gold fluorescence and to determine guidelines for optimal quantification of DNA by SYBR Gold staining, we performed a series of experiments probing the interaction of SYBR Gold with DNA by monitoring bulk fluorescence intensity values. We measured fluorescence intensities as a function of DNA and SYBR Gold concentration using three different techniques: a plate reader, a qPCR cycler with fluorescence intensity readout, and gel electrophoresis (Methods, **Figure 4A-C,** and **Supplementary Figure S3**). The absorbance spectra showed that for concentrations ≤ 1 μM SYBR Gold, the absorbance is ≤ 0.1 (**Figure 4D**), such that inner filter effects (i.e. absorption of the excitation intensity and/or absorption of emitted photons before detection in the measurement cell) could be neglected (36). Fluorescence intensities recorded for a range of DNA and SYBR Gold concentrations ≤ 1 μM increase with increasing DNA concentration for fixed dye concentration (data sets with different color codes in **Figure 4D-F**) and conversely also increased with increasing dye for fixed DNA concentration. The results obtained from the plate reader, the qPCR cycler, and gel electrophoresis are very similar (compare **Figure 4D-F**). The fluorescence data from the three measurement modalities were well described by the finite concentration McGhee-von Hippel model, eqn. (8), with *K*_*d*_, *n*, and *α* as fitting parameters. We find good agreement between the fitted binding parameters for the three different fluorescence measurement modalities and also between the values obtained from single-molecule MT measurements of DNA mechanics and from the fluorescence intensities (**Table 1**). The close agreement between the binding parameters from mechanical manipulation and from fluorescence again strongly suggests that intercalation is the only binding mode of SYBR Gold to DNA that contributes to the fluorescence signal. In addition, the data suggest that the fluorescence intensity is linear in the number of intercalated SYBR Gold molecules, over a broad range of DNA dye molecules per DNA base pair, which in turn rules out significant effects on fluorescence from proximity of SYBR Gold molecules in the helix, e.g. through static quenching effects.

**Table 1.** McGhee-von Hippel model binding parameters determined from different data sets.

| | Dissociation constant<br>$K_d$ ($\mu\text{M}$) | Binding site size<br>$n$ | Fluorescence<br>intensity scale $\alpha$ |
| --- | --- | --- | --- |
| Magnetic tweezers<br>force-extension data | $0.132 \pm 0.013$ | $1.67 \pm 0.04$ | N.A. |
| Plate reader<br>fluorescence | 0.061 | 1.46 | $4.28 \cdot 10^{10}$ |
| qPCR cyclers<br>fluorescence | 0.094 | 1.45 | $2.21 \cdot 10^{10}$ |
| Gel electrophoresis | 0.093 | 1.57 | $7.59 \cdot 10^9$ |

**Figure 4.**
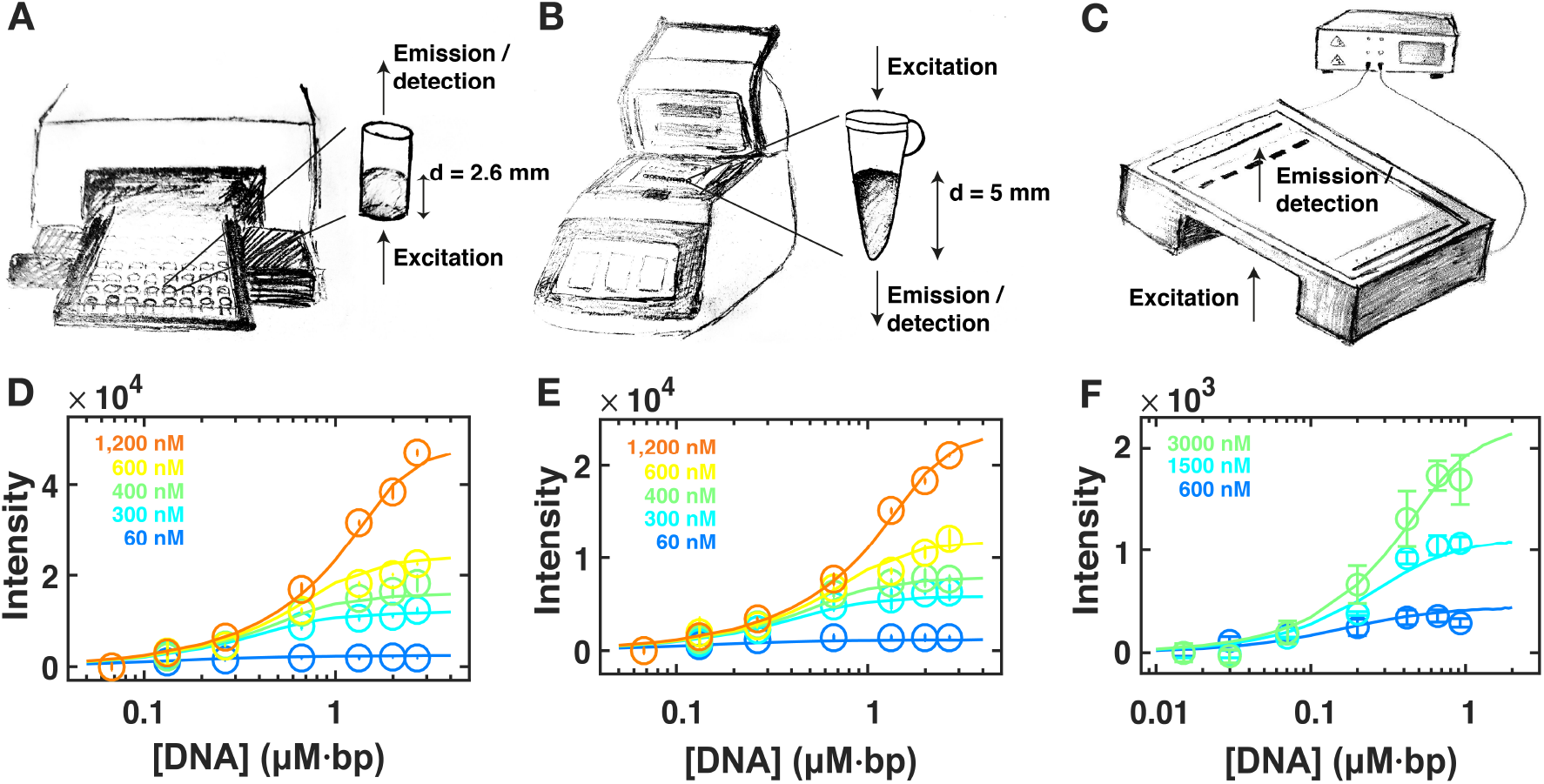
Fluorescence as a function of DNA and SYBR Gold concentrations below 1 μM dye. **A)** Schematic of the 96 well plate reader setup: the SYBR Gold-DNA solution is inserted into the wells of the plate. The sample is excited from the bottom and the fluorescence intensity is measured from the top. The pathlength of the setup is d = (0.26 ± 0.05) mm. **B)** Schematic of the qPCR cycler setup: the SYBR Gold-DNA solution is inserted into PCR tubes that are then placed in the thermal cycler. The sample is excited from the top and the fluorescence intensity is measured from the bottom. The pathlength of the setup is d = (0.50 ± 0.05) mm. **C)** Schematic of the gel electrophoresis setup; the gel is stained with SYBR Gold after running. **D)** Fluorescence intensities recorded using a plate reader and torsionally unstrained DNA (pBR322). The circles and error bars are the mean and standard deviation from at least two independent measurements. The solid line is the best fit of the finite concentration McGhee-von Hippel model (see Methods and **Table 1**). The SYBR Gold concentrations are (from blue to orange) 60, 300, 400, 600, 1200 nM. **E)** Fluorescence intensities recorded using a qPCR cycler. Same conditions and same fitting as for the plate reader data shown in D. **F)** Fluorescence intensities recorded using gel electrophoresis. Same fitting as for the plate reader data shown in D. The SYBR Gold concentrations indicated in the legend text are of the staining solution. The actual concentrations in the gel are lower, but approximately 10-fold, as determined from fitting a global dilution factor (see Methods).

### Dynamic self-quenching limits the linear fluorescence response at high SYBR Gold concentrations

From bulk fluorescence intensity measurements at [*SYBR Gold*] > 1 μ*M*, fluorescence increases with increasing [*SYBR Gold*] were lower than expected (**Figure 5)**. In a first step we used the available absorbance data (**Figure 3A**) to correct for the inner filter effect using the formula (36)

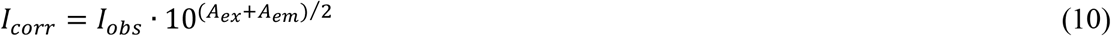

**Figure 5.**
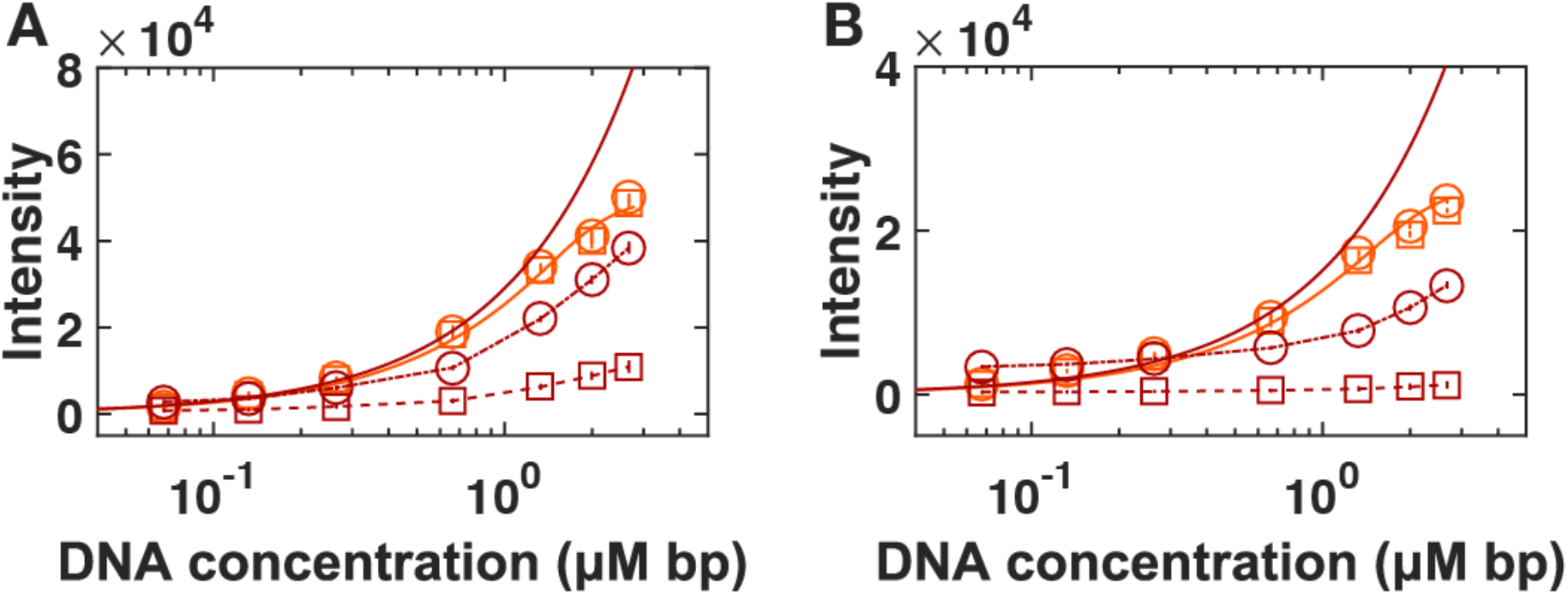
SYBR Gold fluorescence at high dye concentrations. **A)** Fluorescence intensity recorded using a plate reader for torsionally unstrained DNA (pBR322). The orange circles and line are for [SYBR Gold] = 1.2 μM and identical to the data shown in Figure 4C, included as a reference. Red squares are raw data for [SYBR Gold] = 60 μM. Red circles are the same data corrected for the inner filter effect. Dashed and dot-dashed lines are included as guides to the eye. The solid red line is the prediction for [SYBR Gold] = 60 μM using the finite concentration McGhee-von Hippel model and the parameters from Figure 4C. **B)** Fluorescence intensities recorded using a qPCR cycler. Same symbols, conditions, and fitting procedure as for the plate reader data shown in A.

where *I*_*corr*_ and *I*_*obs*_ are the corrected and observed fluorescence intensity and *A*_*ex*_ and *A*_*em*_ the path-length corrected absorbance values. Correcting for the inner filter effect increases the fluorescence values significantly in this concentration regime (**Figure 5**, red circles). Nonetheless, even the corrected fluorescence values are still below the intensities observed at lower dye concentration and much below the values predicted by the finite concentration McGhee-von Hippel model using the parameters determined from the fits to the lower concentration data (**Figure 5**, solid lines). This suggests that some form of self-quenching occurs. The observed quenching above 1 μM SYBR Gold is unlikely due to dye-dye interactions intercalated in the DNA helix, which have been described previously for other dyes (53), as the quenching appears to be similar for different DNA concentration, which correspond to different loading densities in the DNA helix for a given dye concentration.

### Fluorescence lifetime measurements reveal a dynamic quenching at high SYBR Gold concentrations

To better understand the mechanism of self-quenching, we determined fluorescence lifetimes of SYBR Gold by single photon counting. In a first set of experiments we varied the DNA concentration at a constant [*SYBR Gold*] = 1.2 μ*M*. In the absence of DNA, SYBR Gold shows a time-correlated single photon counting histogram that cannot be distinguished from the instrumental response function of our system. DNA binding results in a single exponential fluorescence decay (**Fig 6A,C**) with a lifetime in range of ~6.5 ns. Our lifetime value is similar, but slightly larger than the values reported previously (2). The difference might be due to variations in data analysis (tail fitting vs. reconvolution fitting) and experimental conditions (DNA sample and buffer). At constant [*SYBR Gold*] = 1.2 μ*M*, lifetimes vary only minimally from ~6.9 ns at high loading ratios >1:2 to ~6.1 ns at lower loading density of dye in the DNA helix (<1:10). This is consistent with the absence of any dynamic quenching for [*SYBR Gold*] = 1.2 μ*M*, since dynamic quenching would lead to a reduction in the lifetime with increasing packing density (*i.e*., lower DNA concentration), while we experimentally observe an increase (**Figure 6C**). The slightly longer lifetime at high loading densities is consistent with a stabilization of the excited state and with the observed red shift in the emission at high loading densities (**Figure 3E**), and possibly caused by the stiffening of the helix at high packing densities (**Figure 1D**).

**Figure 6.**
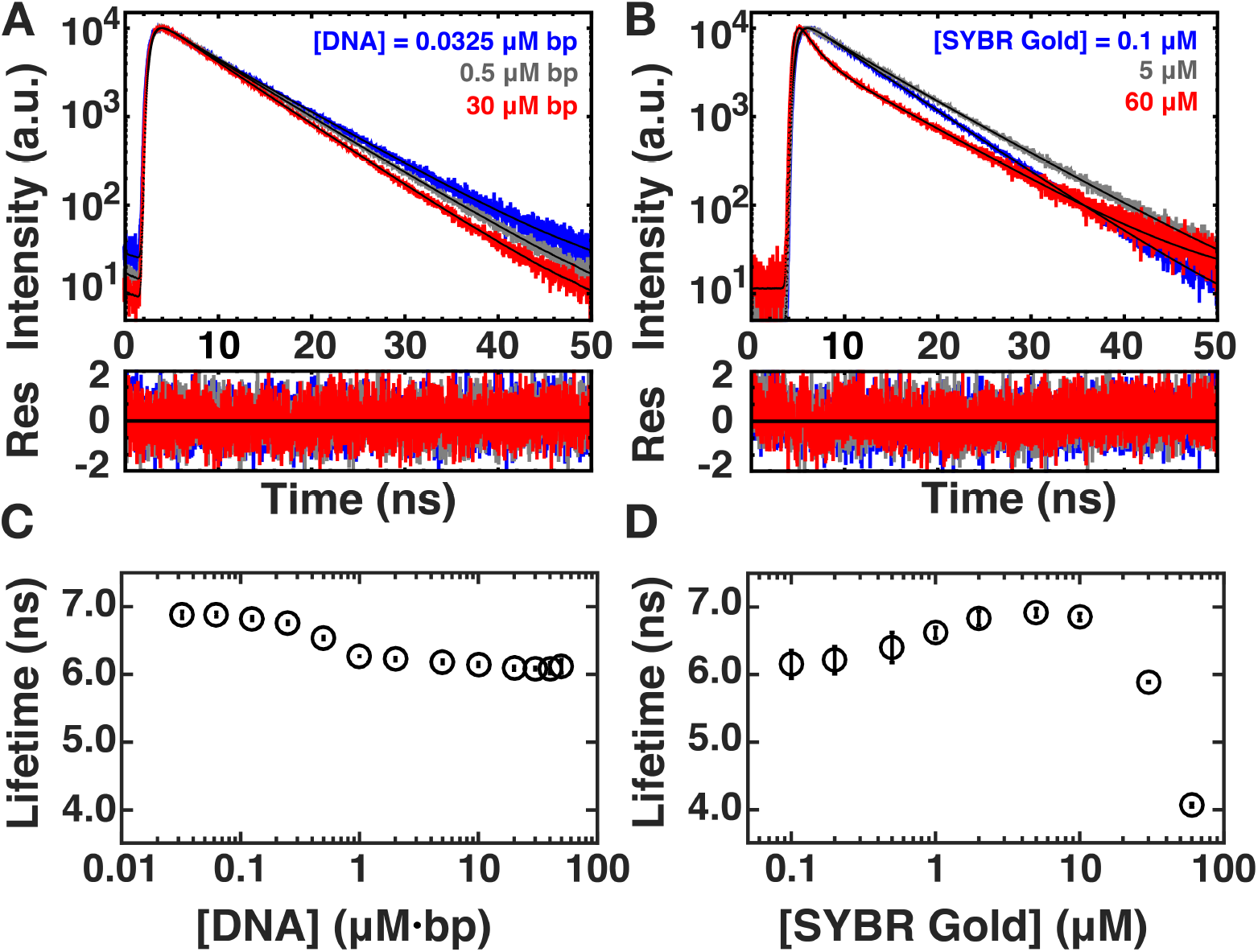
Fluorescence lifetime measurements. **A)** Fluorescence lifetime measurements for 0.0325 (blue), 0.5 (grey) and 30 (red) μM·bp DNA at a SYBR Gold concentration of 1.2 μM. The data are fit by a single exponential decay. **B)** Fluorescence lifetime measurements for 0.1 μM (blue), 5 μM (grey) and 60 μM (red) SYBR Gold in the presence of a constant DNA concentration of 2 μM·bp DNA. At high dye concentrations, the data show a clearly bi-exponential decay, therefore, we fit the data by a two exponential model (Methods). **C)** Fluorescent lifetimes as a function of DNA concentration determined from single-exponential fits as in (A). **D)** Amplitude weighted fluorescent lifetime as a function of SYBR Gold concentration from measurements at 2 μM·bp DNA determined from two exponential fits (Methods).

In a second set of measurements we varied the SYBR Gold concentration while keeping the DNA concentration constant at 2 μM·bp (**Figure 6B** and **6D**). For low [*SYBR Gold*] ≤ 5 μ*M*, fluorescence decays are well described by single exponential fits, and the fitted lifetimes slightly increase up to ~6.9 ns with increasing SYBR Gold concentration (**Figure 6B**). However, at [*SYBR Gold*] > 5 μ*M*, the lifetimes are no longer accurately described by a single exponential fit. Instead, we see a fast decaying component appearing (**Figure 6B**). For consistency, we fit the entire data set with a two-component model, with two exponential decays (Methods). We report an amplitude weighted overall decay constant in **Figure 6D** and the individual decay times and amplitudes in **Supplementary Figure S4**.

The observation of a reduction in lifetime, together with a reduction in fluorescence intensity (**Figure 5**), for [*SYBR Gold*] > 1 μ*M* at constant DNA concentration is indicative of a dynamic self-quenching mechanism. Since the data rule out self-quenching by intercalated dyes in the DNA helix, a more likely scenario is a dynamic quenching mechanism from free or loosely bound dyes in solution. A quenching mechanism from free SYBR Gold in solution would imply for our data at the highest SYBR Gold concentration with a quenching decay time constant of ~1 ns and a free dye concentrations of ~60 μM an apparent bimolecular quenching constant of (~1 ns · 60 μM)^−1^ ~ 10^13^ M^−1^·s^−1^, which is roughly three orders of magnitude larger than diffusion controlled on-rates (~ 10^10^ M^−1^·s^−1^). In addition, quenching from free dye in solution should give rise to a single-exponential decay with a concentration dependent decay constant, while we observe two rate constants. Therefore, the most likely scenario is dynamic self-quenching of the intercalated SYBR Gold from SYBR Gold molecules that are in close proximity to the DNA helix, possibly due to electrostatic interactions (SYBR Gold is positively charged (2) and DNA is highly negatively charged (54)). These charge interactions or, possibly, a combination of charge interactions and other association modes of SYBR Gold with the DNA helix will increase the local effective concentration of SYBR Gold and therefore facilitate dynamic quenching (36).

### Recommendations for quantitation of DNA using SYBR Gold staining

Our single-molecule MT and fluorescence spectroscopy assays provide a comprehensive view of DNA-SYBR Gold interactions. This knowledge eneables us to provids practical guidelines for optimal DNA detection and quantitative DNA sensing applications using SYBR Gold. For quantitative assays, it is desirable to have a linear relationship between fluorescence intensity and DNA concentration. The optimal SYBR Gold concentration to ensure a linear relation between DNA concentration (up to DNA concentrations of ≈ 2 μM·bp or 1.3 ng/μl) and fluorescence intensity as well as an optimal sensitivity for DNA detection is at 1.2 μM (≈1:5000 dilution of the stock solution). This value for the optimal SYBR Gold concentration is 2x larger than the manufacturer’s recommendation of 1:10000 fold dilution. Reliable detection is possible with lower dye concentrations, however, with a reduced range for a linear fluorescence-DNA concentration response (**Figure 7A)**. So if linearity and high signal are important, using high dye concentrations is desirable. However, at very high dye concentration ([*SYBR Gold*] > 1.2 μM quenching and inner filter effects become relevant and need to be corrected for to get accurate and quantitative measuring results (**Figure 7A** and **Fig 7B**). For many measurements, it is likely beneficial to avoid inner filter and quenching effects by keeping [*SYBR Gold*] ≤ 1.2 μM.

**Figure 7.**
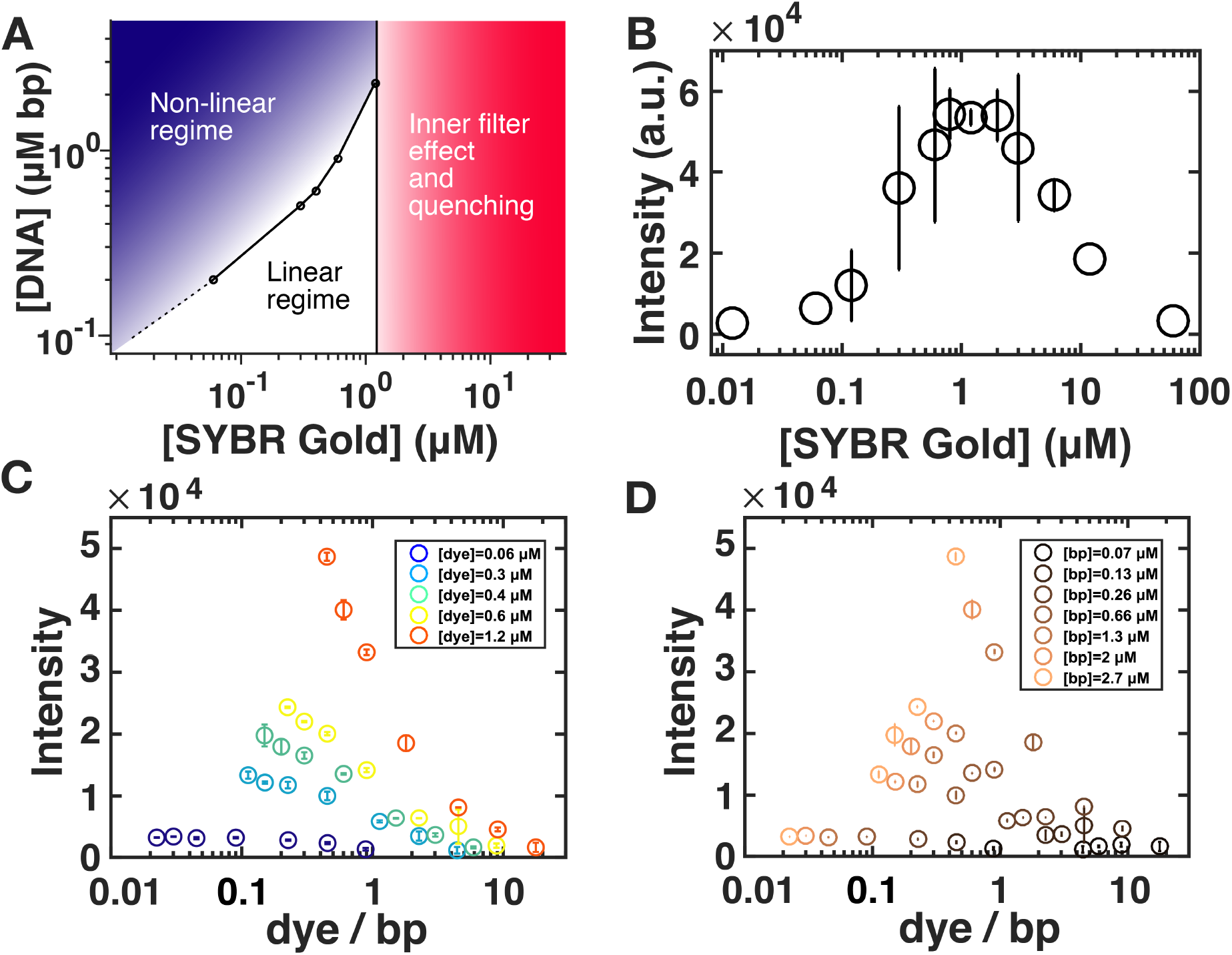
Recommendations for DNA quantitation by SYBR Gold fluorescence. **A)** Phase diagram depicting different regimes of DNA detection by SYBR Gold fluorescence, as a function of SYBR Gold and DNA concentrations. **B)** Fluorescence intensity recorded using a qPCR cycler at constant DNA concentration (Lambda DNA, 2.7 μM bp) at varying SYBR Gold concentrations. The data points are the mean values of two independent experiments including their standard deviations or stem from just one experiment. **C)** Fluorescence intensity recorded using a plate reader as a function of the dye per base ratio. The circles and error bars are the mean and standard deviation from at least two independent measurements. The SYBR Gold concentrations are (from blue to orange) 0.06, 0.3, 0.4, 0.6, 1.2 μM. These are the same data as in Figure 4D plotted as a function of dyes per bp. **D)** Fluorescence intensity recorded using a plate reader as a function of the dye per base ratio. The circles and error bars are the mean and standard deviation from at least two independent measurements. The DNA base pair concentrations are (from dark to light brown) 0.07, 0.13, 0.26, 0.66, 1.3, 2, 2.7 μM.

In addition, we note that that the fluorescence intensity depends, in general, on both the dye and DNA concentration. Reporting concentrations as “dye per base pair” –as is frequently done for DNA stains– is problematic, since a large range of intensities correspond to the same dye per base pair ratio (**Figure 7C,D**) and since the connection between dye to base pair ratio and fluorescence intensity is many-to-one. Therefore, we advise to report both DNA and dye concentrations explicitly.

## CONCLUSIONS

We have employed single-molecule MT that provide rotational control in addition to control of the stretching forces in order to probe the binding properties of SYBR Gold to DNA. The single-molecule MT assay reveals systematic lengthening (up to 1.7 times the DNA contour length in the absence of dye) and unwinding of DNA (19.1° ± 0.7° per SYBR Gold bound) upon SYBR Gold binding. The mechanical signature is fully consistent with intercalation, with an unwinding angle at the low end of the range of previously investigated intercalators. Fitting the McGhee-von Hippel model to the MT data, we find a binding constant *K*_*d*_ = 0.132 ± 0.013 μM and binding site size of *n* = 1.67 ± 0.04. These findings are in good agreement with the parameters from fluorescence intensity experiments for dye concentrations of up to ~1.2 μM suggesting that the intercalative binding mode is responsible for the observed fluorescence. Additionally, we find that for SYBR Gold concentrations > 1.2 μM, fluorescence quenching and inner filter effects become relevant. While we see no evidence for quenching between dyes intercalated in the helix, our data provide clear evidence for dynamic quenching from dyes in a loosely bound (like at least partially driven by electrostatics) „cloud“ around the DNA. Overall, we found that while SYBR Gold is advantageous due to its high quantum yield and brightness, it has a relatively narrow range of concentrations that strike a balance between avoiding inner filter effects and quenching, while staining DNA with a linear fluorescence to DNA concentration relationship. In summary, our work shows how using complementary techniques can provide a highly quantitative and comprehensive view of DNA-small molecule interactions and we anticipate our approach to be broadly applicable to other DNA binding agents.

## ACKNOWLEDGEMENTS

We thank Alex Urban and Carola Lampe for help with recording excitation and emission spectra, Dieter Braun and Patrick Kudella for help with qPCR cycler measurements, and Aidin Lak for useful discussions.

## FUNDING

Funded by the Deutsche Forschungsgemeinschaft (DFG, German Research Foundation) though SFB 863 – Project ID 111166240 (to J.L.), SFB863 – Project A13 (to T.C), GRK2062 – project C03 (to T.C.), and the European Commission (ERC starting grant ERC-StG 638536 SM-IMPORT to T.C.). C.G. acknowledges a PhD fellowship from the Studienstiftung des deutschen Volkes.

## Notes

### Competing Interest Statement

The authors have declared no competing interest.

## REFERENCES

1. Tuma, R.S., Beaudet, M.P., Jin, X., Jones, L.J., Cheung, C.Y., Yue, S. and Singer, V.L. (1999) Characterization of SYBR Gold nucleic acid gel stain: a dye optimized for use with 300-nm ultraviolet transilluminators. Anal Biochem, 268, 278–288.

2. Cosa, G., Focsaneanu, K.S., McLean, J.R.N., McNamee, J.P. and Scaiano, J.C. (2001) Photophysical Properties of Fluorescent DNA-dyes Bound to Single- and Double-stranded DNA in Aqueous Buffered Solution¶. Photochemistry and photobiology, 73, 585–599.

3. Biebricher, A.S., Heller, I., Roijmans, R.F., Hoekstra, T.P., Peterman, E.J. and Wuite, G.J. (2015) The impact of DNA intercalators on DNA and DNA-processing enzymes elucidated through force-dependent binding kinetics. Nat Commun, 6, 7304.

4. McKillip, J.L. and Drake, M. (2004) Real-Time Nucleic Acid–Based Detection Methods for Pathogenic Bacteria in Food. Journal of Food Protection, 67, 823–832.

5. Uslan, C. and Şebnem Sesalan, B. (2013) The synthesis, photochemical and biological properties of new silicon phthalocyanines. Inorganica Chimica Acta, 394, 353–362.

6. Kuhler, P., Roller, E.M., Schreiber, R., Liedl, T., Lohmuller, T. and Feldmann, J. (2014) Plasmonic DNA-origami nanoantennas for surface-enhanced Raman spectroscopy. Nano Lett, 14, 2914–2919.

7. Wang, J.C. (1974) The degree of unwinding of the DNA helix by ethidium. I. Titration of twisted PM2 DNA molecules in alkaline cesium chloride density gradients. J Mol Biol, 89, 783–801.

8. Langner, K.M., Kedzierski, P., Sokalski, W.A. and Leszczynski, J. (2006) Physical nature of ethidium and proflavine interactions with nucleic acid bases in the intercalation plane. The journal of physical chemistry, 110, 9720–9727.

9. Cluzel, P., Lebrun, A., Heller, C., Lavery, R., Viovy, J.L., Chatenay, D. and Caron, F. (1996) DNA: an extensible molecule. Science, 271, 792–794.

10. Bennink, M.L., Scharer, O.D., Kanaar, R., Sakata-Sogawa, K., Schins, J.M., Kanger, J.S., de Grooth, B.G. and Greve, J. (1999) Single-molecule manipulation of double-stranded DNA using optical tweezers: interaction studies of DNA with RecA and YOYO-1. Cytometry, 36, 200–208.

11. Tessmer, I.a.B., C. G. and Skinner, G. M. and Molloy, J. E. and Hoggett, J. G. and Tendler, S. J. B. and Allen S. (2003) Mode of drug binding to DNA determined by optical tweezers force spectroscopy. J. Mod. Opt., 50, 1627–1636.

12. Sischka, A., Toensing, K., Eckel, R., Wilking, S.D., Sewald, N., Ros, R. and Anselmetti, D. (2005) Molecular mechanisms and kinetics between DNA and DNA binding ligands. Biophys J, 88, 404–411.

13. Mihailovic, A.a.V., I. and McCauley, M. and Ly, E. and Williams, M.C. and Spain, E.M. and Nu–ez, M.E. (2006) Exploring the interaction of ruthenium(II) polypyridyl complexes with DNA using single-molecule techniques. Langmuir, 22, 4699––4709.

14. Vladescu, I.D., McCauley, M.J., Rouzina, I. and Williams, M.C. (2005) Mapping the phase diagram of single DNA molecule force-induced melting in the presence of ethidium. Phys Rev Lett, 95, 158102.

15. Vladescu, I.D., McCauley, M.J., Nunez, M.E., Rouzina, I. and Williams, M.C. (2007) Quantifying force-dependent and zero-force DNA intercalation by single-molecule stretching. Nat Methods, 4, 517–522.

16. Paramanathan, T., Westerlund, F., McCauley, M.J., Rouzina, I., Lincoln, P. and Williams, M.C. (2008) Mechanically manipulating the DNA threading intercalation rate. Journal of the American Chemical Society, 130, 3752–+.

17. Yang, T.S., Cui, Y., Wu, C.M., Lo, J.M., Chiang, C.S., Shu, W.Y., Chung, W.J., Yu, C.S., Chiang, K.N. and Hsu, I.C. (2009) Determining the zero-force binding energetics of an intercalated DNA complex by a single-molecule approach. Chemphyschem, 10, 2791–2794.

18. Paik, D.H. and Perkins, T.T. (2012) Dynamics and multiple stable binding modes of DNA intercalators revealed by single-molecule force spectroscopy. Angew Chem Int Ed Engl, 51, 1811–1815.

19. Schakenraad, K., Biebricher, A.S., Sebregts, M., Ten Bensel, B., Peterman, E.J.G., Wuite, G.J.L., Heller, I., Storm, C. and van der Schoot, P. (2017) Hyperstretching DNA. Nat Commun, 8, 2197.

20. Manosas, M., Camunas-Soler, J., Croquette, V. and Ritort, F. (2017) Single molecule high-throughput footprinting of small and large DNA ligands. Nat Commun, 8, 304.

21. Clark, A.G., Naufer, M.N., Westerlund, F., Lincoln, P., Rouzina, I., Paramanathan, T. and Williams, M.C. (2018) Reshaping the Energy Landscape Transforms the Mechanism and Binding Kinetics of DNA Threading Intercalation. Biochemistry, 57, 614–619.

22. Krautbauer, R., Pope, L.H., Schrader, T.E., Allen, S. and Gaub, H.E. (2002) Discriminating small molecule DNA binding modes by single molecule force spectroscopy. FEBS Lett, 510, 154–158.

23. Eckel, R., Ros, R., Ros, A., Wilking, S.D., Sewald, N. and Anselmetti, D. (2003) Identification of binding mechanisms in single molecule-DNA complexes. Biophys J, 85, 1968–1973.

24. Lipfert, J., Klijnhout, S. and Dekker, N.H. (2010) Torsional sensing of small-molecule binding using magnetic tweezers. Nucleic Acids Res, 38, 7122–7132.

25. Salerno, D., Brogioli, D., Cassina, V., Turchi, D., Beretta, G.L., Seruggia, D., Ziano, R., Zunino, F. and Mantegazza, F. (2010) Magnetic tweezers measurements of the nanomechanical properties of DNA in the presence of drugs. Nucleic Acids Res, 38, 7089–7099.

26. Gunther, K., Mertig, M. and Seidel, R. (2010) Mechanical and structural properties of YOYO-1 complexed DNA. Nucleic Acids Res, 38, 6526–6532.

27. Wang, Y., Sischka, A., Walhorn, V., Tönsing, K. and Anselmetti, D. (2016) Nanomechanics of Fluorescent DNA Dyes on DNA Investigated by Magnetic Tweezers. Biophysical Journal, 111, 1604–1611.

28. Kriegel, F., Ermann, N., Forbes, R., Dulin, D., Dekker, N.H. and Lipfert, J. (2017) Probing the salt dependence of the torsional stiffness of DNA by multiplexed magnetic torque tweezers. Nucleic Acids Res, 45, 5920–5929.

29. Lipfert, J., Hao, X. and Dekker, N.H. (2009) Quantitative modeling and optimization of magnetic tweezers. Biophys J, 96, 5040–5049.

30. Kriegel, F., Ermann, N. and Lipfert, J. (2017) Probing the mechanical properties, conformational changes, and interactions of nucleic acids with magnetic tweezers. J Struct Biol, 197, 26–36.

31. Cnossen, J.P., Dulin, D. and Dekker, N.H. (2014) An optimized software framework for real-time, high-throughput tracking of spherical beads. Rev Sci Instrum, 85, 103712.

32. te Velthuis, A., Kerssemakers, J.W.J., Lipfert, J. and Dekker, N.H. (2010) Quantitative Guidelines for Force Calibration through Spectral Analysis of Magnetic Tweezers Data. Biophysical Journal, 99, 1292–1302.

33. Lansdorp, B.M. and Saleh, O.A. (2012) Power spectrum and Allan variance methods for calibrating single-molecule video-tracking instruments. Rev Sci Instrum, 83, 025115.

34. Bouchiat, C., Wang, M.D., Allemand, J., Strick, T., Block, S.M. and Croquette, V. (1999) Estimating the persistence length of a worm-like chain molecule from force-extension measurements. Biophysical Journal, 76, 409–413.

35. van Oene, M.M., Dickinson, L.E., Pedaci, F., Kober, M., Dulin, D., Lipfert, J. and Dekker, N.H. (2015) Biological magnetometry: torque on superparamagnetic beads in magnetic fields. Phys Rev Lett, 114, 218301.

36. Lakowicz, J.R. (2013) Principles of fluorescence spectroscopy. Springer Science & Business Media.

37. McGhee, J.D. and Hippel, P.H.v. (1974) Theoretical Aspects of DNA-Protein Interactions :Co-operative and Non-co-operative Binding of Large Ligands to a One-dimensional Homogeneous Lattice. J. Mol. Biol., 86, 469–489.

38. Lerman, L.S. (1961) Structural considerations in the interaction of DNA and acridines. J Mol Biol, 3, 18–30.

39. Biebricher, A.S., Heller, I., Roijmans, R.F.H., Hoekstra, T.P., Peterman, E.J.G. and Wuite, G.J.L. (2015) The impact of DNA intercalators on DNA and DNA-processing enzymes elucidated through force-dependent binding kinetics. Nature Communications, 6, 7304.

40. Bustamante, C., Marko, J.F., Siggia, E.D. and Smith, S. (1994) Entropic elasticity of lambda-phage DNA. Science, 265, 1599–1600.

41. Press, W.H., Flannery, B.P., Teukolsky, S.A. and Vetterling, W.T. (1992) Numerical Recipes in C, Cambridge University Press. New York.

42. Berman, H.M. and Young, P.R. (1981) The interaction of intercalating drugs with nucleic acids. Annu Rev Biophys Bioeng, 10, 87–114.

43. Wang, Y., Schellenberg, H., Walhorn, V., Toensing, K. and Anselmetti, D. (2017) Binding mechanism of PicoGreen to DNA characterized by magnetic tweezers and fluorescence spectroscopy. European Biophysics Journal, 46, 561–566.

44. Vanderlinden, W., Kolbeck, P.J., Frederickx, W., Konrad, S.F., Nicolaus, T., Lampe, C., Urban, A.S., Moucheron, C. and Lipfert, J. (2019) Ru(TAP)32+ uses multivalent binding to accelerate and constrain photo-adduct formation on DNA. Chem Commun (Camb), 55, 8764–8767.

45. Quake, S.R., Babcock, H. and Chu, S. (1997) The dynamics of partially extended single molecules of DNA. Nature, 388, 151–154.

46. Bordelon, J.A., Feierabend, K.J., Siddiqui, S.A., Wright, L.L. and Petty, J.T. (2002) Viscometry and Atomic Force Microscopy Studies of the Interactions of a Dimeric Cyanine Dye with DNA. The Journal of Physical Chemistry B, 106, 4838–4843.

47. Dikic, J. and Seidel, R. (2019) Anticooperative Binding Governs the Mechanics of Ethidium-Complexed DNA. Biophys J, 116, 1394–1405.

48. Clauvelin, N., Audoly, B. and Neukirch, S. (2008) Mechanical response of plectonemic DNA: An analytical solution. Macromolecules, 41, 4479–4483.

49. Vilfan, I.D., Lipfert, J., Koster, D.A., Lemay, S.G. and Dekker, N.H. (2009) In Hinterdorfer, P. and van Oijen, A. (eds.), Handbook of Single-Molecule Biophysics. Springer.

50. Neukirch, S. and Marko, J.F. (2011) Analytical description of extension, torque, and supercoiling radius of a stretched twisted DNA. Phys Rev Lett, 106, 138104.

51. Strick, T.R., Dessinges, M.N., Charvin, G., Dekker, N.H., Allemand, J.F., Bensimon, D. and Croquette, V. (2003) Stretching of macromolecules and proteins. Rep. Prog. Phys., 66, 1–45.

52. Celedon, A., Wirtz, D. and Sun, S. (2010) Torsional mechanics of DNA are regulated by small-molecule intercalation. The journal of physical chemistry, 114, 16929–16935.

53. Benvin, A.L., Creeger, Y., Fisher, G.W., Ballou, B., Waggoner, A.S. and Armitage, B.A. (2007) Fluorescent DNA Nanotags: Supramolecular Fluorescent Labels Based on Intercalating Dye Arrays Assembled on Nanostructured DNA Templates. Journal of the American Chemical Society, 129, 2025–2034.

54. Lipfert, J., Doniach, S., Das, R. and Herschlag, D. (2014) Understanding nucleic acid-ion interactions. Annu Rev Biochem, 83, 813–841.

