## Supplementary Information for "Intercalative DNA binding governs fluorescence enhancement of SYBR Gold"

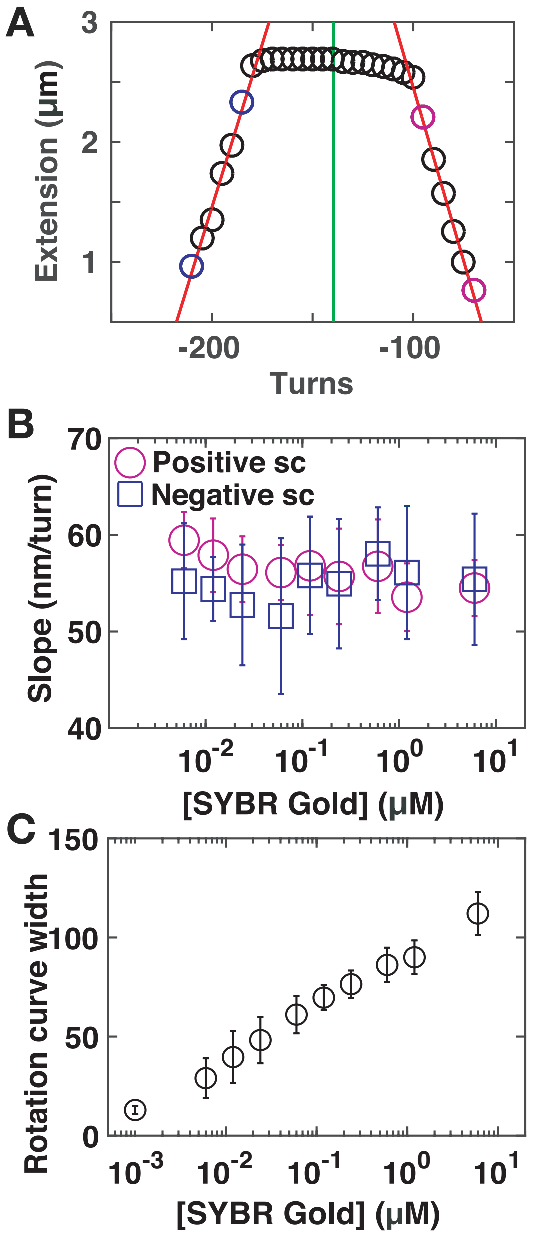


**Supplementary Figure S1. SYBR Gold rotation curve analysis. A)** Example of a rotation-extension curve and analysis of center positions and slopes. Data are for 7.9 kb DNA at $F$ = 0.5 pN in the presence of 240 nM SYBR Gold (circles). The center positions are determined from fitting slopes in the positive and negative plectonemic regime (red lines) and by computing the intersection of the two slopes (indicated by the green line). **B)** Extension vs. turn slopes in the plectonemic regime (determined as indicated by the red lines in panel A) as a function of SYBR Gold concentration for positive (red circles) and negative (blue squares) plectonemic supercoils. **C)** Width of the pre-buckling regime (in turns) vs. SYBR Gold concentration. Data points and error bars in panel B and C are the mean and standard deviation from at least 14 independent measurements. In panel A one typical experiment is shown for clarity.

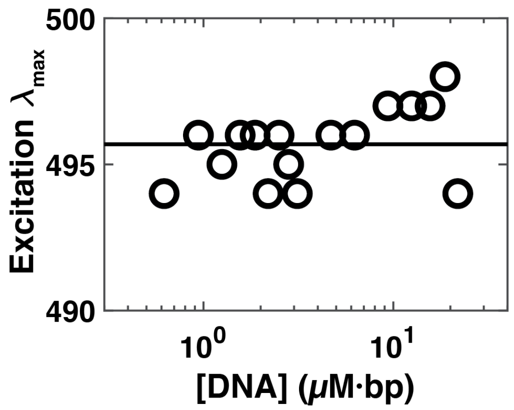


**Supplementary Figure S2. Position of the excitation maxima at constant SYBR Gold concentration (1.2 µM) and varying DNA concentrations**. No significant shift is observed for the excitation maxima. The line is simply the mean of the data points at all concentrations 495.7 nm ± 1.25 nm (mean ± std).

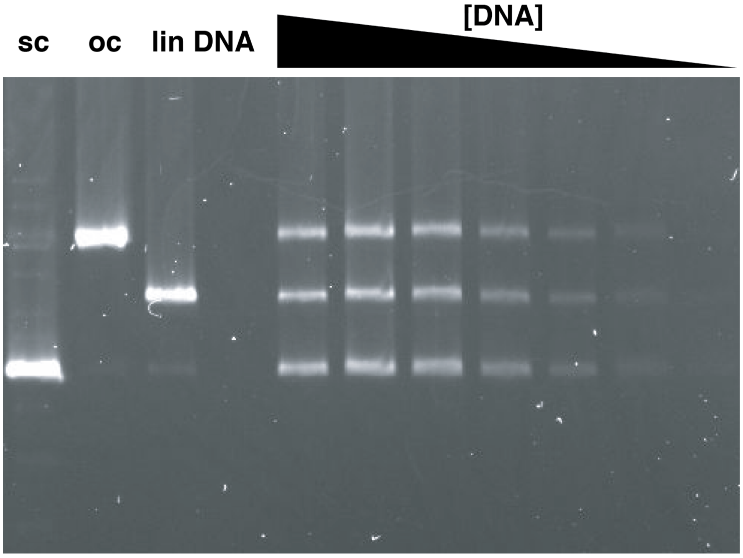


**Supplementary Figure S3. Gel electrophoresis of PBR322 plasmid DNA and visualization by SYBR Gold fluorescence.** Lanes from left to right: supercoiled DNA, open-circular DNA, linear DNA, empty lane, and equimolar mixtures of supercoiled, open-circular, and linear DNA with decreasing amounts of DNA: 30 ng, 21.4 ng, 13.6 ng, 6.5 ng, 2.3 ng, 1.0 ng, 0.5 ng. The gel was stained after electrophoretic separation with 1.5 μM SYBR Gold in TAE buffer (see Methods).

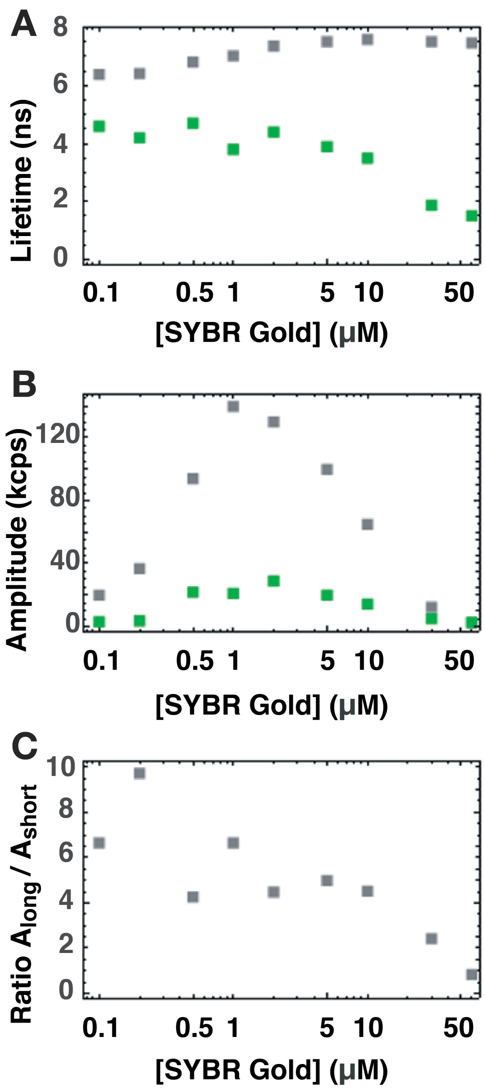


**Supplementary Figure S4. Fluorescence lifetime measurements. A)** Fluorescent lifetimes as a function of SYBR Gold concentration at a constant concentration of 2 µM·bp DNA determined from bi-exponential fits. These are the same data as shown in Figure 6B and D. The fit gives a shorter lifetime (green, ~2-4 ns) that decreases strongly with increasing SYBR Gold concentration and a longer lifetime (gray, ~6-7 ns) that slightly increases with increasing SYBR Gold concentration. **B)** Amplitudes of the two fluorescence lifetimes vs. SYBR Gold concentration; longer lifetime in green, shorter lifetime in grey. **C)** Ratio of the two amplitudes of the fluorescence lifetimes vs. SYBR Gold concentration. Up to concentrations of ~5 µM SYBR Gold, the long lifetimes dominates the fit, with the short lifetime contributing < 25% of the total amplitude. At higher concentration, both amplitudes decrease as the total intensity strongly decreases, but the shorter lifetime –which we attribute to dynamic quenching from SYBR Gold molecules close to the DNA– becomes more important and roughly equal in magnitude to the long component.
